# METLIN Neutral Loss Database Enhances Similarity Analysis

**DOI:** 10.1101/2021.04.02.438066

**Authors:** Aries Aisporna, H. Paul Benton, Jean Marie Galano, Martin Giera, Gary Siuzdak

**Affiliations:** Scripps Center for Metabolomics and Mass Spectrometry, The Scripps Research Institute, 10550 North Torrey Pines Road, La Jolla, CA 92037; Institut des Biomolécules Max Mousseron, UMR 5247 CNRS, ENSCM, Université de Montpellier, France; Leiden University Medical Center, Center for Proteomics and Metabolomics, Albinusdreef 2, 2333ZA Leiden, Netherlands; Department of Chemistry, Molecular and Computational Biology, The Scripps Research Institute, 10550 North Torrey Pines Road, La Jolla, CA 92037

## Abstract

Tandem mass spectrometry (MS^2^) data is an effective resource for the identification of known molecules and the putative identification of novel, previously uncharacterized molecules (unknowns). Yet, MS^2^ data alone is limited in characterizing structurally closely related molecules with different masses. Neutral loss data is key in retrieving this structural similarity. To facilitate unknown identification and complement METLIN’s MS^2^ fragment ion data for characterizing structurally related molecules, we have created the METLIN neutral loss database (https://metlin-nl.scripps.edu).

---

Similarity analysis^1–4^ and molecular networking^5,6^ using tandem mass spectrometry (MS^2^) data have become valuable approaches for identifying previously uncharacterized molecules (unknowns).^1^ Yet key structural information can be lost when relying solely on this fragment ion data, for example, the loss of a sulfate ion from two similar molecules of different masses will not result in fragment ion overlap.^7^ This is of significant practical relevance. A user who would try to identify an unknown based on a database similarity search would not succeed in obtaining structurally useful matches. However, retrieving this structurally useful information is possible by analyzing the differences between the molecular ion and the fragment ion, or better known as the neutral loss (Δ*m/z*). Neutral losses^1,2^ constitute a rich resource, and have already been widely used in proteomics, pharmacology, and metabolomics for over three decades.^1,2,8-12^ Yet, even though mass spectrometry-based neutral loss (NL) analysis has been extensively applied, with hundreds to thousands of papers on the topic, no comprehensive small molecule library of neutral loss data exists.

The new METLIN neutral loss database (METLIN-NL) has been created from METLIN’s 850,000 MS^2^ small molecule molecular standards database to facilitate neutral loss searching. The neutral loss data was derived across a broad range of standards representing hundreds of different chemical classes.^3,13^ METLIN’s MS^2^ data was converted to METLIN-NL spectra (e.g. **Figure 1** asymmetric dimethylarginine (ADMA)) by calculating the differences between the precursor molecular ion and the fragment ions in the experimental MS^2^ mass spectra (**Figure 1a**). The neutral loss spectra (NL_intensity_ vs Δ*m/z*) were created (e.g. ADMA **Figure 1b**) with the neutral loss intensity (NL_intensity_) using the fragment ion intensities from each precursor/fragment generated neutral loss (Δ*m/z*). It should be noted that not all precursor to fragment peaks represent a true neutral loss between the precursor and fragment ions, and therefore some of the peaks in the NL spectra can also be considered (as recently described) hypothetical neutral losses.

**Figure 1.**
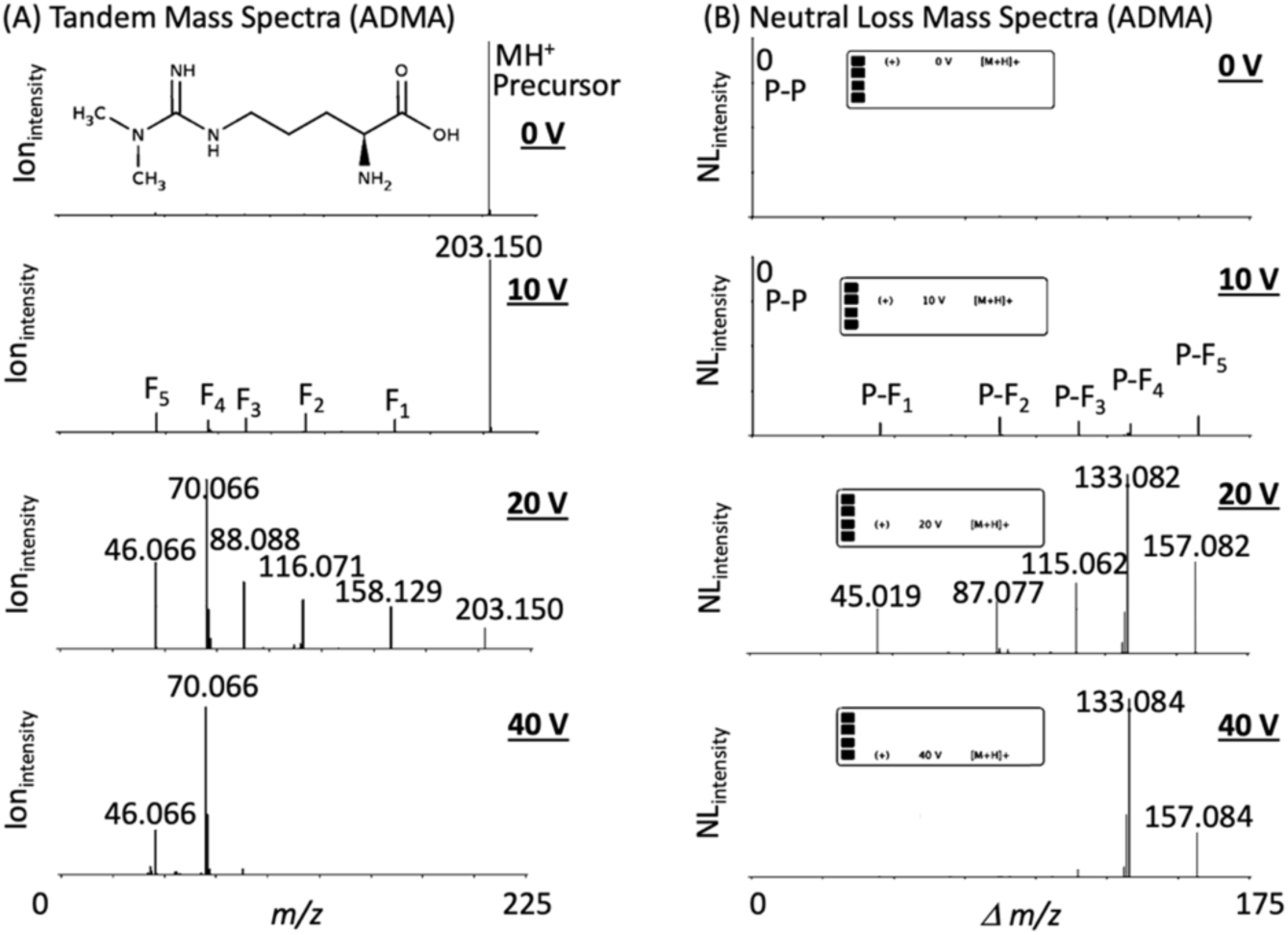
The METLIN-NL mass spectral database was derived from the METLIN MS^2^ data on over 850,000 molecular standards, and their respective fragment ions. (A) Asymmetric dimethylarginine (ADMA) and its representative METLIN tandem mass spectra at four different collision energies. (B) METLIN-NL spectra (NL_intensity_ vs. Δm/z) of ADMA was generated by calculating the difference between the precursor and fragment ions with NL_intensity_ based on the original fragment ion intensities. “P” refers to precursor ion and “F” refers to fragment ion.

METLIN-NL is a compilation NL_intensity_ vs Δ*m/z* spectra generated from METLIN’s eight distinct MS^2^ data sets^3^ created from 850,000 standards. This compilation is represented within METLIN-NL at four different collision energies and in both positive and negative ionization modes. The rationale behind providing multiple conditions is that MS^2^ collision energies have not been standardized and such broad acquisition parameters are required to represent the output across different instrument types. An additional rationale for the array of conditions is that different molecules can fragment differently depending on the collision energies thus METLIN provides a broad range of empirical data across its 850,000 standards. It is worth noting that all of METLIN’s MS^2^ data is empirical data and has not been generated using predictive *in silico-based* approaches.

A secondary set of METLIN-NL data has also been accumulated based on precursor minus fragment ion transitions as well as all possible fragment to fragment ion transitions to provide a more comprehensive set of experimentally derived structural data. Unlike the original METLIN MS^2^ database, METLIN-NL represents a translation that more effectively enables the molecular annotation of unknown molecular entities since NL data is inherently corrected for molecular weight differences (**Figures 2**).

**Figure 2.**
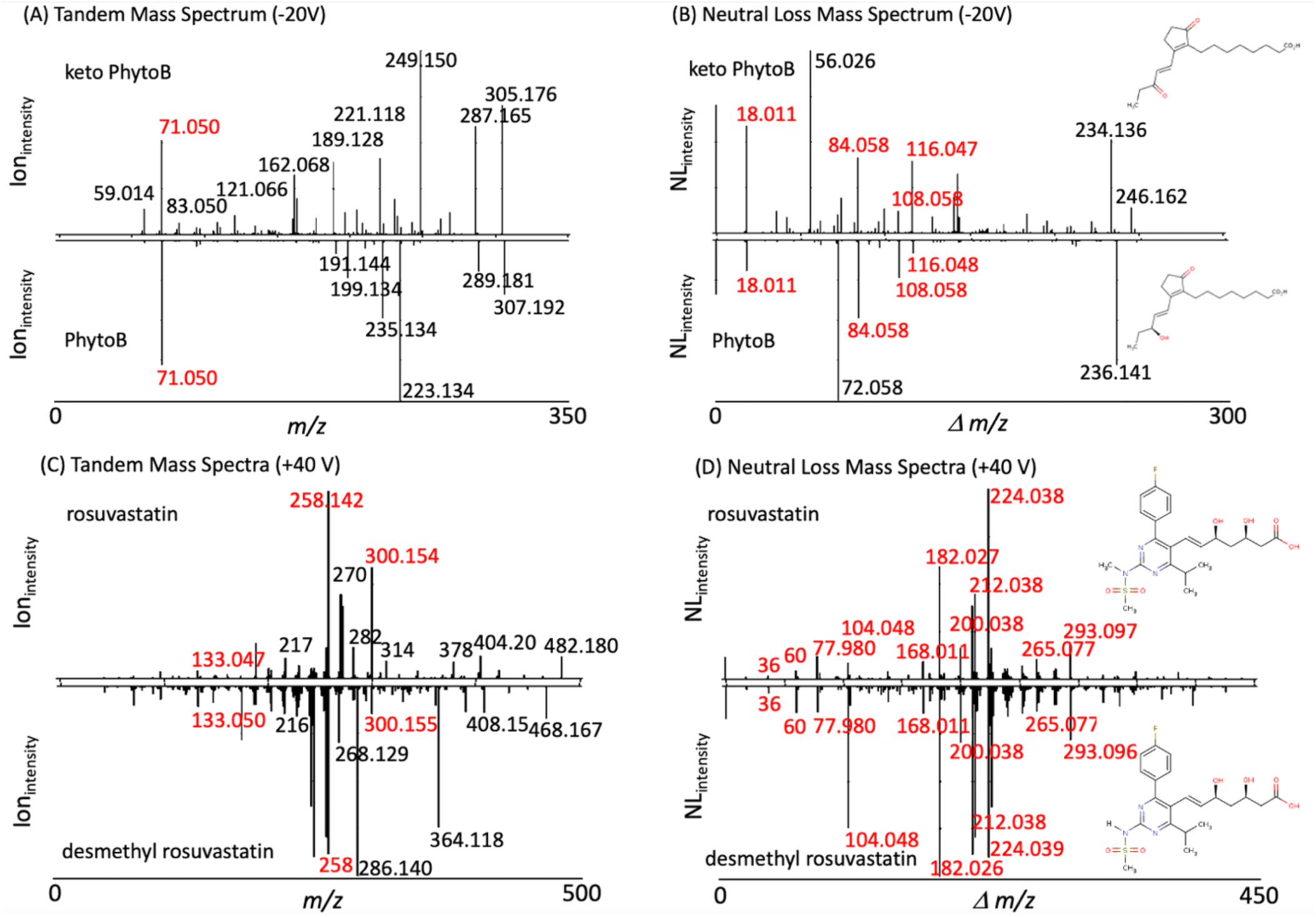
MS^2^ and neutral loss data on two related oxylipins (16 keto 16-B_1_-PhytoP and 16-B_1_-PhytoP) and the statin drugs rosuvastatin and desmethyl rosuvastatin. (A) Oxylipin MS^2^ data show little overlap (in red) in contrast to the (B) neutral loss spectra with the high resolution neutral loss data facilitating similarity analysis with both providing complementary structural information. (C) MS^2^ and (D) neutral loss data on rosuvastatin and desmethyl rosuvastatin. MS^2^ data show few overlapping peaks (in red) while the neutral loss spectra provide near complete overlap. Interestingly, while the neutral losses help facilitate similarity, the MS^2^ data provides more structural information on their structurally distinguishing features.

To test the utility of METLIN-NL we examined two different types of molecular structures, oxylipins and a pharmaceutical (statin) drug and its demethylated metabolite. Oxylipins^14^ represent a class of highly active lipid metabolites ubiquitous in humans and plants, and specifically, the phytoprostanes (PhytoPs) class of oxylipins resemble prostaglandin-like compounds that are found in seeds and vegetable oils derived from oxidative cyclization of α-linolenic acid. Since PhytoPs are a class of highly structurally related oxylipins and are suspected to have additional unidentified analogs,^14–16^ we chose them to demonstrate the utility of METLIN-NL. Tandem MS and neutral loss data were recently generated on a set of PhytoPs, including the structural analogs 16-B_1_-PhytoP and 16-keto 16-B_1_-PhytoP (**Figure 2**). When trying to extrapolate/correlate the observed tandem MS spectra of the two PhytoPs, classic similarity searching was of very limited value providing only one overlapping ion, even though some fragments presented an expected two Dalton difference (**Figure 2a**). This exemplifies that two structurally very similar molecules can yield highly different MS^2^ spectra limiting similarity searching possibilities and thereby severely impacting the usefulness of this approach for the identification of chemically closely related substances. However neutral loss similarity analysis yielded multiple overlapping neutral losses (**Figure 2b**). Further analysis of the tandem MS data as well as the molecular weight difference between the two molecules being 2 Daltons, were consistent with 16-keto 16-B1-PhytoP. This neutral loss data (unlike the MS^2^ data) helped to easily correlate the two molecules, and the distinguishing neutral losses and fragment ions exclusive to 16-keto 16-B_1_-PhytoP and 16-B_1_-Phyto provides significant structural information.

The purpose of having a large database is to help reduce the need for speculation, and allow for the rapid identification of molecules. However, since many molecular structures are not represented in any database, similarity analyses offer an alternative in the preliminary characterization process. This process extends beyond naturally occurring molecules and can be applied just as readily to xenobiotics and other chemical entities. The second example in applying METLIN-NL is shown here for a non endogenous drug molecule and its metabolite.

The well known cholesterol-lowering statin drug rosuvastatin^17^ (trade name Crestor) and its active metabolite desmethyl rosuvastatin^18^ differ in mass by 14 Daltons (demethylation reaction) and the MS^2^ and neutral loss data (**Figure 2c & 2d**) of these two molecules have recently been acquired and populated within METLIN and METLIN-NL. As was observed with the oxylipins, tandem MS data was of limited utility when searching METLIN (**Figure 2c**), where 3 fragment ions were overlapping between the two molecules. However neutral loss matching/detection showed near complete overlap (**Figure 2d**). Further analysis of the tandem MS data as well as the molecular weight difference between the two molecules being 14 Daltons, were consistent with loss of a methyl group. For the rosuvastatin NL data, the overlap in the neutral loss data clearly dominated the comparative analyses, making similarity searching much more effective using neutral loss while the MS^2^ data provided complementary information that was informative for structural determination. Overall, the neutral loss data which was completely derived from the MS^2^ data, is more effective (than MS^2^) at showing similarity.

METLIN’s molecular standards with systematically acquired experimental MS^2^ data across multiple collision energies, allows for the comprehensive generation and graphical user interface (*beta*) visualization (**Figure 3**) of neutral loss data. Fragment ion and neutral loss similarity analysis^1^ was originally developed to aid in the identification of novel molecules (unknowns)^1^ by using fragment ion and neutral loss data to help align an unknown molecule to compounds with similar fragmentation data within a database. However now, with a neutral loss database of small molecules via METLIN-NL, neutral loss similarity analysis can be more readily applied to a host of biological and chemical challenges.

**Figure 3.**
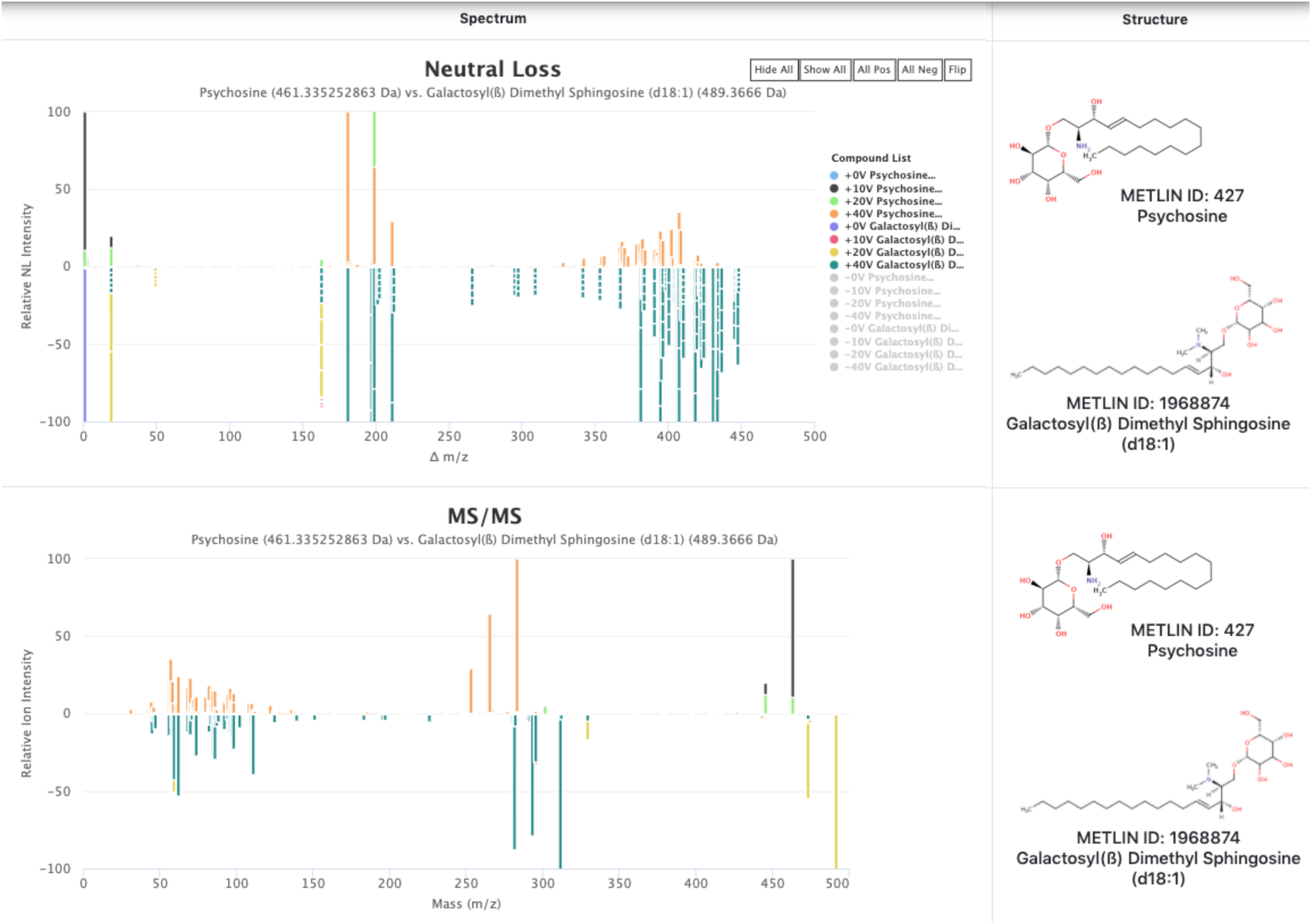
METLIN-NL is built on a Linux platform with this beta version of the graphical user interface (GUI) created using Highcharts, HTML, JQuery and PHP. The beta GUI allows for comparative analyses between different compounds including neutral loss data (NL_int_ vs Δm/z) as well as MS/MS data (Frag_int_ vs m/z) in both positive and negative ionization modes and either at each individual collision energy, or a composite of multiple collision energies, as shown here for psychosine and gal dimethyl sphingosine. The Neutral Loss and MS/MS spectra are a composite of all the collision energies in positive ionization mode.

Overall, METLIN-NL empirically derived data will enable new types of analyses facilitating more rapid identification of unknown compounds via both fragment ion and neutral loss similarity searching.^2^ Both biologists and chemists will be able applying METLIN-NL to the structure elucidation of unknowns derived from animals,^19^ plants,^14,20^ or microbiota^21^; and METLIN-NL can also be used as a resource for identifying unexpected synthetic chemical or enzymatically modified drug products (e.g. pharmaceuticals^22^) as it is populated with both biological and chemical entities. Given METLIN’s extensive userbase,^3^ and the ubiquitous application of mass spectrometry-based neutral loss analysis (dating back three decades), METLIN-NL promises to have wide-ranging utility.

## Acknowledgements

This research was partially funded by National Institutes of Health grants R35 GM130385 (G.S.), P30 MH062261 (G.S.), P01 DA026146 (G.S.), and U01 CA235493 (G.S.) and by Ecosystems and Networks Integrated with Genes and Molecular Assemblies (ENIGMA), a Scientific Focus Area Program at Lawrence Berkeley National Laboratory for the U.S. Department of Energy, Office of Science, Office of Biological and Environmental Research, under contract number DE-AC02-05CH11231 (G.S.).

## AUTHOR CONTRIBUTIONS

A.A., H.P.B., J.M.G., M.G. and G.S. contributed to data collection, analysis, and manuscript writing.

## Notes

https://metlin-nl.scripps.edu

### Competing Interest Statement

The authors have declared no competing interest.

### Summary of Updates

Improved resolution on Figure 1 correcting an error the chemical structure.

